# Bird song comparison using deep learning trained from avian perceptual judgments

**DOI:** 10.1101/2022.12.23.521425

**Authors:** Lies Zandberg, Veronica Morfi, Julia George, David F. Clayton, Dan Stowell, Robert F. Lachlan

## Abstract

Our understanding of bird song, a model system for animal communication and the neurobiology of learning, depends critically on making reliable, validated comparisons between the complex multidimensional syllables that are used in songs. However, most assessments of song similarity are based on human inspection of spectrograms, or computational methods developed from human intuitions. Using a novel automated operant conditioning system, we collected a large corpus of zebra finches’ (*Taeniopygia guttata*) decisions about song syllable similarity. We use this dataset to compare and externally validate similarity algorithms in widely-used publicly available software (Raven, Sound Analysis Pro, Luscinia). Although these methods all perform better than chance, they do not closely emulate the avian assessments. We then introduce a novel deep learning method that can produce perceptual similarity judgements trained on such avian decisions. We find that this new method outperforms the established methods in accuracy and more closely approaches the avian assessments. Inconsistent (hence ambiguous) decisions are a common occurrence in animal behavioural data; we show that a modification of the deep learning training that accommodates these leads to the strongest performance. We argue this approach is the best way to validate methods to compare song similarity, that our dataset can be used to validate novel methods, and that the general approach can easily be extended to other species.

## Introduction

Bird song is an important model system for several related fields: the neurobiology of learning, animal communication, sexual selection and cultural evolution (Catchpole & Slater, 2003; Whiten, 2019; Bolhuis *et al*., 2010). Similarities in development, neurobiology and neurogenomics have further led song to become a model system for understanding human speech (Bolhuis *et al*., 2010; ten Cate *et al*., 2013). Its importance largely results from the fact that songbirds memorise songs they hear from adult conspecifics during a sensitive phase early in life, and then produce imitations of these songs (Catchpole & Slater, 2003), but this flexibility also leads to bird song being an unusually variable animal signal.

All these fields of bird song research depend on reliable comparisons of song recordings. This task is not straightforward because bird songs, like many acoustic signals, are very high dimensional—with many potential spectral features varying dynamically through the course of each song unit. In fact, when animals themselves compare two vocal signals, they must subjectively integrate numerous differences in frequency and timing. This leads to the realisation that the comparison methods we use need to be validated against animals’ own perception: there is no comparison method that can be objectively “correct” (Janik, 1999; ten Cate *et al*., 2013).

The first studies comparing bird song were performed by comparing songs by ear or transcribing songs into onomatopoetic descriptions (Marler, 1952). The invention of the sonograph revolutionized the study of animal communication (Thorpe, 1954), making it possible to produce a visual representation of the time-frequency structure of the songs on basis of which spectrograms of songs can be compared visually: human visual assessment of spectrographic similarity (HVA). To assess similarity more repeatably, Clark *et al*. (1987) proposed spectrographic cross correlation (SPCC) as a computational measure of similarity. Two spectrograms are superimposed and then shifted temporally to find the peak correlation coefficient, which is used as a measure of similarity between the sounds. A recent implementation of this method can be found in software such as Raven Pro (Center for Conservation Bioacoustics, 2019). SPCC relies heavily on the spectrographic representation of the signal, for example being intolerant of often minor differences in the relative duration of spectral components. An alternative approach is to extract contours of acoustic features that are believed to be perceptually relevant, and then to align the contours of two signals and integrate the differences between them in these features. Sound Analysis Pro (SAP) is a widely used tool that employs this approach, measuring several features (e.g. Wiener entropy, spectral continuity, pitch and frequency modulation), that can be tuned to the study species (Tchernichovski et *al*., 2000). SAP allows for timing differences by linearly warping time by up to 30%. Another software package, Luscinia (Lachlan, 2020), also uses multiple acoustic features to generate a dissimilarity measure, but uses dynamic time-warping (DTW) to align features. DTW allows non-linear warping of time to find the optimal alignment of features. SAP and DTW both require decisions about which acoustic features to use and how to weight them.

Recently, data-driven methods have been proposed for analysis of song similarity based on machine learning (Mets & Brainard, 2018; Goffinet *et al*., 2021; Sethi *et al*., 2020). While not yet widely used in bird song research, such methods reduce the dependence on engineering good features, by using spectrograms or waveforms directly as input. Deep learning can achieve impressive accuracy on various tasks, but this neither implies nor demands that their recognition strategies are similar to those of animals (Fel et al., 2022). All comparison methods make assumptions about what constitutes similarity in acoustic signals. This is true even in the case of unsupervised deep learning, which (like principal components analysis before it) derives a representation automatically from unlabelled data: in such a case, assumptions about similarity emerge implicitly from the choice of training data as well as the neural net structure.

The most fundamental problem facing all comparison methods is how their assumptions are validated (Janik, 1999). The design of the algorithms relies on the intuition of the scientists, although particular decisions can be justified on the basis of perceptual research. But, typically, they have only been validated by comparison with the previous “gold standard”, HVA. However, HVA is based on human, not avian, perception, in the visual, rather than auditory domain, of a spectrographic representation of a sound, which differs in known ways from how sounds are perceived (e.g. the linear rather than logarithmic scaling of frequency (Dooling & Prior, 2017a)).

To overcome these problems, in the present work we have: 1) developed a novel method to test zebra finches for their perception of sound similarity and deployed it to collect a large data-set of avian sound assessments, 2) developed a novel algorithm to train a deep neural network to assess song similarity, and 3) validated this algorithm as well as current conventional methods of measuring song similarity on basis of the birds’ assessments. By using an automated operant feeder in combination with radio-frequency identification (RFID) technology we were able to individually train and test group-living zebra finches continuously, enabling us to collect over 900,000 responses to stimuli. Such response data typically involve some amount of inconsistent or unclear responses, so we further adapted our algorithm to train using both the unambiguous and the ambiguous operant response data. Although here we tested birds for their assessments of similarity in zebra finch syllables, this method can be applied to measure similarity between arbitrary new song unit recordings. Unlike previous methods, this algorithm is not validated by HVA, but by a measure of the birds’ own perceptions of sound similarity. Here we found that the algorithm we developed on basis of the birds’ assessments outperformed all widely used methods.

## Methods

### Assessments of song similarity

#### Birds

We experimentally tested group-living zebra finches for their perception of sound similarity using a novel operant conditioning system. Experiments were performed using 26 domesticated zebra finches from an outbred colony at Queen Mary University of London. All birds used in the experiment were hatched in the population between December 2015 and May 2018 and reared in a flock in a large free-flight room. The birds were fitted with a RFID (radio frequency identification) tag leg band (Eccel Technology Ltd., Leicester, UK) in addition to the standard metal leg ring and a colour band. Throughout the experiment all birds were kept at a 12.00:12.00 light:dark schedule (lights on at 7:00 GMT), in two separate aviary rooms (one room with two aviaries of approx. 100×200×200cm and one room with two aviaries of approx. 200×200×200cm). Each experimental aviary held 4-6 individuals in both single sex and mixed groups. The birds received water *ad libitum*, and a commercial tropical seed mixture from the operant feeders. Each aviary was outfitted with 2 operant feeders. Birds could feed *ad libitum* for a half hour after lights on, and a half hour before lights off. Operant feeders were operational from 7:30 to 18:30, which meant that the birds had to interact with the feeder to gain access to the food.

#### Operant device

We used two different operant feeder designs (see Figure 1a for design 1). Each feeder device consisted of a PIT tag detection system, a motor activated feeding tube, two capacitive touch perches registering responses, speaker system, a Raspberry Pi 3B+ and a Hall effect sensor. In design 1 all electronics are integrated in a weatherproof electronics box, whereas in design 2 these components are integrated in drainpipe elements. Additionally, in design 2 the stimuli are played through two speakers, whereas in design 1 there is only one speaker. Other than the external casing and the number of speakers the devices work identically. The Raspberry Pi computer receives information from the PIT tag system and the response perches, and on basis of this input controls the speaker output and the running of the motor activating the reward mechanism. A PIT antenna was positioned at 20cm from the feeding tube, which registered the bird passing through. On opposite sides of the feeding tube two aluminium response perches were placed, which registered the presence of a bird on the perch through a capacitive touch sensor. The feeding mechanism consisted of a PVC outer tube with an opening on both the left and the right side, and inside this tube a transparent acrylic inner tube with only one opening. This inner tube held all the seeds used as a reward. By turning the inner tube so that the opening lined up with the opening in the outer tube, birds were able to access the seeds in the inner tube. To prevent misalignment of the inner tube a Hall effect sensor was built in together with a magnet attached to the inner tube, to check the position of the inner tube after each reward, and to readjust when needed.

**Figure 1:**
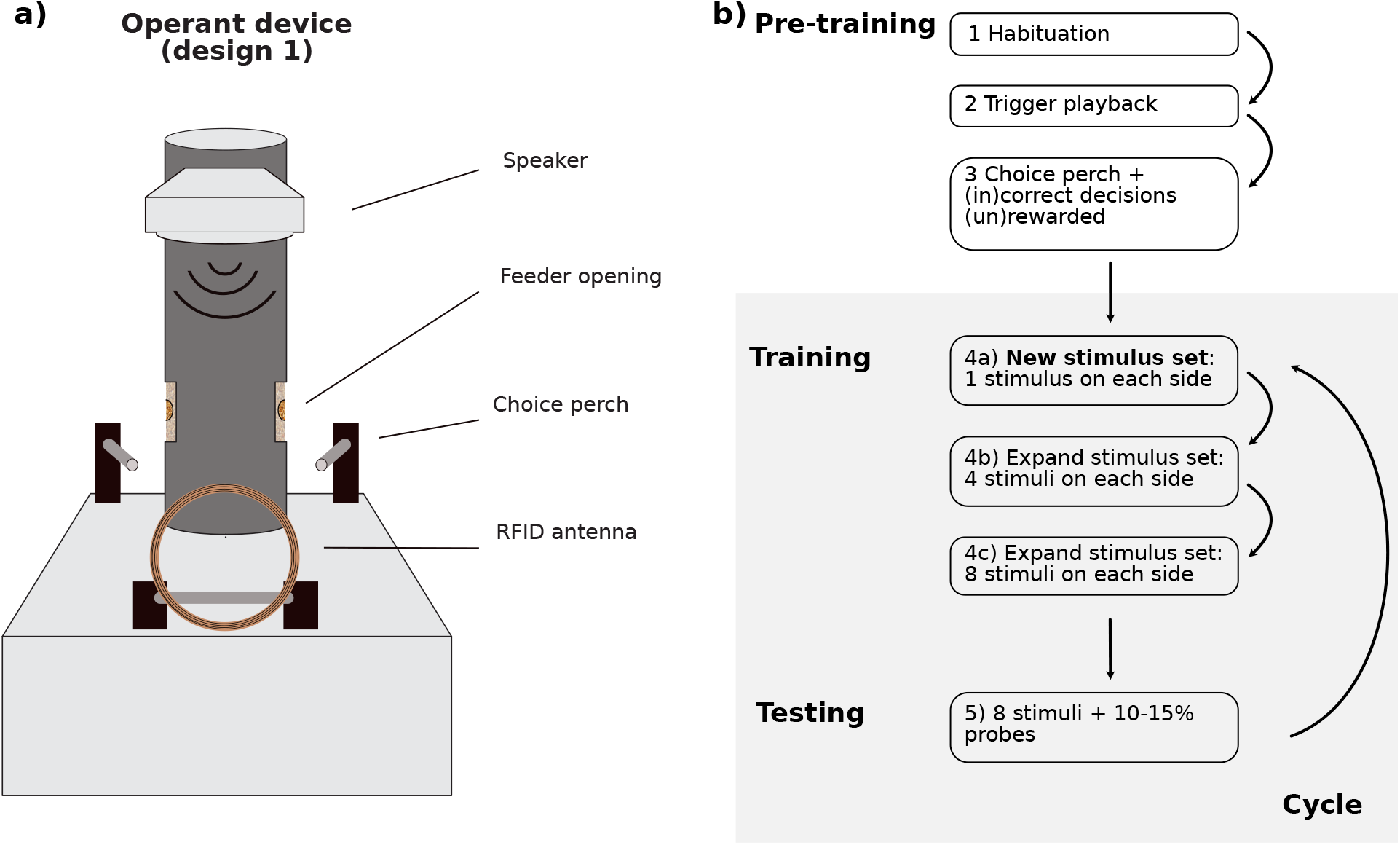
**a)** Schematic image of operant feeder design 1. **b)** Bird training scheme with the different training steps.

#### Sound stimuli

Sound stimuli used were 887 syllables extracted from zebra finch song recordings (Boogert *et al*., 2018). These song recordings were made in a different population from the focal population to avoid any potential confounding effects in case of recognition of singer identity. All songs were recorded at 48kHz and divided into separate syllables, which were high pass filtered (100Hz), normalised and a 20ms fade in and out was added. When a stimulus was played it was repeated 4 times in quick succession.

#### Test paradigm

We tested group-living captive zebra finches for their perception of sound similarity using an operant feeder with a 2 alternative forced choice (2AFC) paradigm: forced-choice ‘AXB’ judgments (see Figure 1a). In short, this meant that we trained birds to discriminate between two sound stimuli, ‘A’ and ‘B’ and associate each of them with one of the choice perches. Once the birds were accurate in their responses to these training stimuli we introduced 10–15% probe stimuli ‘X’, which in terms of similarity lies somewhere in between the two sets of training stimuli, for them to assess for their perception of similarity. When a bird was presented with a probe stimulus we expected it to respond by choosing the perch of the training stimulus that it perceived to be most similar to the presented probe. We infer that the choice made in response to the AXB (probe-stimulus) triplet represents a perception of categorical similarity. A trial was started by a bird passing through the PIT antenna and registering its identity on basis of the PIT tag. Depending on the identity of the bird, the software selected a stimulus sound to be played through the speaker(s) on the top of the device. Each stimulus sound is either a training stimulus, and thus associated with a ‘correct’ side of the device, left or right, or is a probe sound. Responses to training sounds were rewarded only when the bird responded by hopping onto the perch associated with that side. In case of an incorrect response, a short fragment of white noise was played and the feeder remained closed. All responses to probe sounds were rewarded by the inner feeding tube opening, through which the birds had access to the seeds for 2 seconds. For a more detailed description of the steps involved in pre-training, training and probe presentation see below and Figure 1b).

#### Selection of stimuli

Using Luscinia sound software (Lachlan, 2020) we compared all syllables with each other and extracted similarity measures for each combination of syllables. On basis of these similarity measures, we subsequently constructed a matrix with, for each syllable, a ranking of all other syllables ranging from the most to the least similar syllable. From all syllables a random syllable was chosen as the first training stimulus, stimulus ‘A’, and a second training stimulus, stimulus ‘B’, was selected to be ranked between 50 and 150 similar to the first training stimulus. For each stimulus A and B, their 7 most similar syllables were selected as additional stimuli. Probe stimuli were selected to be ranked between 20 and 200 of both training stimuli (see Figure S1).

#### Pre-training

*Step 1* – Habituation to the feeder: Feeder opening moves from left to right side, every 30 minutes. Birds can feed *ad libitum* from the feeder. *Step 2* - Birds have to go through the antenna, which triggers a playback of a stimulus. After the playback the feeder immediately opens to the side of the feeder associated with the sound. The feeder remains open for 8 seconds during which the bird can feed. *Step 3* - The time delay to open after playback is increased. When a bird hops onto a choice perch within the time delay period it receives either a correct (stimulus is repeated and the feeder opens immediately) or an incorrect response (playback of 3s white noise, and an added time delay of 30s before the next trial can be initiated). If the bird does not make a choice within the time delay period the feeder opens automatically after the time delay. Step-by-step the time delay period is increased to encourage the birds to make a choice, rather than to wait for the feeder to open automatically. At the same time, the reward time of the automatic opening is reduced from 8 seconds to 3 seconds, whereas the choice-reward time is kept at 8 seconds.

#### Training

*Step 4a* – The time delay to open automatically is removed and from this phase onward the feeder does not open automatically anymore: the birds have to make a correct choice to receive a food reward and the the choice-reward time is reduced to 2 seconds. Similar to the previous phase an incorrect response by the bird is followed by a playback of 3s white noise, and an added time delay of 30s before the next trial can be initiated. In this training phase each bird is trained with its own set of stimuli: one stimulus is rewarded on the left side, one stimulus on the right side. *Step 4b* - When on average the group makes 70% accurate decision we increased the number of stimuli on each side to 4. These stimuli are selected to be the original stimuli plus the 3 syllables most similar to each of the original stimuli. *Step 4c* - When on average the group makes 70% accurate decision we increased to 8 stimuli on each side (original stimulus and the 7 most similar stimuli).

#### Probe testing

*Step 5* - After the group reaches 70% accurate decisions with 8 stimuli on each side we add 10-15% probes. The responses to probe sounds are always rewarded.

#### Assessing birds’ assessments

To assess whether birds made choices we expected, we compared the birds’ assessments of similarity with assessments made by Luscinia. For each stimulus we ranked all other stimuli on basis or their similarity, and determined the rank of the each probe stimulus. For each stimuli-probe triplet we calculated the probe rank difference between the left and the right stimulus: a negative rank difference suggests that, according to Luscinia, the probe is more similar (closer in rank) to the left stimulus, and a positive rank difference suggests that the probe stimulus is more similar to the right stimulus. To account for training stimuli that were closer together or further apart we adjusted the rank difference by dividing it by the mean rank difference. To test whether the birds choices were more similar to Luscinia than expected by chance we compared the birds’ decisions to the distribution of random choice by randomising their decisions 10,000 times. We calculated this for birds with a cycle accuracy of > 65% (overall accuracy in its responses to the training stimuli in a test period).

### Deep learning embeddings

We next built a deep learning model that can learn from the birds’ judgements about similarity of sounds and use it to create a perceptual space that can mirror birds’ perception. This perceptual space is a learnt vector representation of data referred to as an *embedding* space and can be used for classification, verification and other similarity-based tasks (Bromley *et al*., 1993; Bredin, 2017; Sethi *et al*., 2020). Our deep learning approach follows that of Morfı *et al*. (2021), but is adapted to the specific situation in which with animal behavioural data there can be a high proportion of uncertain (ambiguous) data points, due to the limited accuracy and consistency of measured animal judgments. In common with Kumari *et al*. (2019), our method is distinct from most deep learning embedding methods (including triplet-based methods) in that no semantic “classification” labels are used. It is also distinct from purely unsupervised methods which are guided only by the signal variation in the dataset, and thus have no necessary link to perceptual or communicative relevance (Goffinet *et al*., 2021).

In order to infer a data-driven similarity algorithm from our probe responses, it was required to fit a highly non-linear function to the data, where the algorithm input is the audio data, and the fit is conditioned on the responses to our probes. Deep learning has made great advances in creating learnt *embeddings* by methods such as triplet networks which rely on distance metrics rather than classification as the training objective (Schultz & Joachims, 2004; Weinberger & Saul, 2009; Wang *et al*., 2014; Hoffer & Ailon, 2015). Triplet networks are trained on triplets of data points consisting of an anchor, a positive sample, and a negative sample. Through the training procedure an embedding space is learnt so that the anchor and the positive samples are featured close to each other, while at the same time the anchor and negative samples are separated as much as possible. However, most previous successes in representation learning are driven by datasets with explicit class labels, which are used to provide a strong signal of semantic distance even for triplet networks (Thakur *et al*., 2019). In other words, the distance metric is often derived from an underlying measure of classification correctness, rather than general similarity.

Our operant AXB experiments were designed so a probe X can correspond to a triplet anchor. Based on the A/B decision a bird made for that probe, all training stimuli rewarded on the selected side correspond to positive exemplars for that anchor, while all training stimuli rewarded on the opposite side correspond to negative exemplars in the triplet. This makes it possible for us to create a dataset of triplets out of the decisions made by the zebra finches in order to train a triplet network to learn an embedding space for the recordings used in the experiment. We are able to then project new stimuli into this learnt embedding space to measure their similarity to the training stimuli. This is done without any use of labels or class knowledge for the stimuli. The goal is for the learnt embedding space to make the same A/B decisions as birds would make for the recordings.

#### Network architecture

An overview of the deep learning model architecture is depicted in Figure S2. We compute the log scaled power mel spectrogram from the zebra finch recordings, of 150 mel bands and 170 time frames, and use it as input to our model. The selection of log mel spectrogram as input was based on prior work which found these useful for deep learning applied to birdsong data ((Morfı *et al*., 2021)). The architecture of our model consists of a shared convolutional part followed by two parallel parts of attention pooling and max pooling, the predictions of which are combined at the last layers and projected to an embedding space of *d* dimensions (see Morfı *et al*. (2021) for more information).

For our triplet model, the anchor, positive and negative examples are each passed separately through this network, to estimate the coordinates of each stimulus in the perceptual embedding space. The distances between these coordinates are then used as input to the triplet loss function, described next, which will be used to update the weights of the network.

We reiterate that, in common with Kumari *et al*. (2019), our method is distinct from most deep learning embedding methods (including triplet-based methods) in that no semantic “classification” labels are used. Our algorithm training is driven by the data coming from two-alternative similarity decisions, using both the ambiguous and unambiguous behavioural data.

#### Metric learning with triplet loss

Let 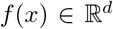 represent an embedding that maps a sample *x* (an audio clip) into a *d*-dimensional space. Often *f*(*x*) is normalised to have unit length for training stability (Schroff *et al*., 2015). We want to ensure that an anchor sample *x_a_* is closer to positive samples *x_p_* that are more perceptually similar to it than negative samples *x_n_* without having any information about the class of each sample. We define the distance between any two points *x_i_* and *x_j_* as *D_ij_*. In our experiments, this distance denotes the squared the Euclidean distance. For a triplet 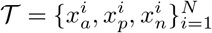 of example stimuli the loss is given by

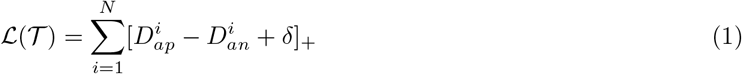

where [·]_+_ is standard hinge loss and *δ* is a non-negative margin hyper-parameter. The larger the margin, the further the negative example should be from the anchor in the embedding space (Hoffer & Ailon, 2015; Bredin, 2017; Thakur *et al*., 2019).

#### Unambiguous triplet setting

In order to build a system that can model the way birds perceive sounds, we require a set of easily distinguishable signals to avoid perceptual ambiguity, which in turn requires a way to estimate the perceptual similarity of such signals. Bird decisions for sound similarity can contain a lot of noise; this can be due to the cycle accuracy of the individual, scrounging interactions or the consistency of side decisions for probes caused by perceptual ambiguity of sounds. In triplet learning it can be important to create a dataset without any ambiguity in it, and in our experiments we try to limit this noise from the above sources as much as possible.

In order to reduce decision noise and make our dataset unambiguous we perform the following pre-processing steps on the bird decision data:

- Discard individual’s decisions if cycle accuracy for them is less than 65% on the training stimuli. (We use training stimuli to estimate this, since we cannot calculate it from the probes: we do not make any assumptions about their positive/negative association with the training stimuli.)
- Discard decisions if there were any technical issues with the device for that day.
- Discard inconsistent probe decisions: a consistent probe decision is one where the same side was chosen for that probe during a cycle by an individual over 70% of the times it was played to them.

This unambiguous dataset of decisions formulates 2,099 triplets. We split these into training and evaluation sets of 1,239 and 860 triplets, respectively. The training triplets are used as input to our deep learning model in order for it to learn a space that can satisfy the triplet metric of Equation 1. The evaluation set only consists of triplets acquired from birds with accuracy of 77% and higher to make it as unambiguous as possible and to properly evaluate the performance of our model.

#### Ambiguous triplet setting

In Kumari *et al*. (2019) a method is introduced to learn perceptual embeddings for haptic signals, gathered from human participants. Similarly to our data, theirs was based on two-alternative decisions about texture similarity between triplets of items. Their model is trained on both unambiguous and ambiguous triplets and they show an improvement over training only on unambiguous data.

To make use of the ambiguous data points for training our model, we collect bird decisions that had inconsistency in side selection. Decisions with 50% to 70% side consistency were chosen from individuals with cycle accuracy of 65% and higher to produce a dataset of “ambiguous” triplets. This ambiguous dataset of decisions totals 1,073 triplets. We split these into training and evaluation sets of 668 and 405 triplets, respectively. Unlike common triplet-based learning approaches which tend to ignore these type of triplets as uninformative, our approach treats both triplet types (ambiguous and unambiguous) as informative, based on the work in Kumari *et al*. (2019).

Depending on whether a triplet is ambiguous or unambiguous, a different condition needs to be satisfied. For an unambiguous triplet 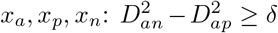; on the other hand for an ambiguous triplet 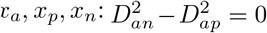, where *D_ij_* is the Euclidean distance between any two points *x_i_* and *x_j_* and *δ* is a non-negative margin hyper-parameter. These conditions are defined as:

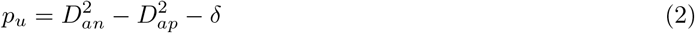

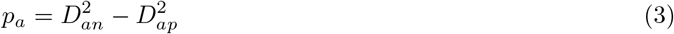

In order to facilitate training with both ambiguous and unambiguous triplets, for a triplet 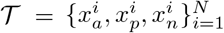 we adjust the loss function of Equation 1 to:

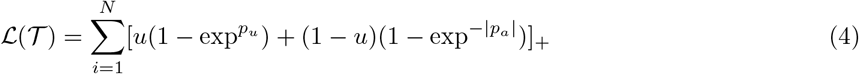

where [·]_+_ is standard hinge loss and *u* ∈ [0,1] denotes if a triplet is ambiguous (*u* = 0) or unambiguous (*u* = 1).

In some of our machine learning tests we use only the unambiguous decisions; we also test the use of ambiguous and unambiguous data together. Furthermore, we explore models that use triplets created by Luscinia after tuning the algorithms parameters (as explained in the following section). In either case the training data are treated as a single training dataset, to train the network by backpropagation using the Adam optimiser, following standard practice in deep learning.

#### Evaluating deep learning

To explore the sensitivity of our deep learning method to hyperparameter settings, we evaluated its performance at different dimensionalities of embedding space, and different ways of using bird decisions and Luscinia-U decisions in training (Table 1). To evaluate whether an algorithm can produce similarity measures compatible with the birds’ judgments recorded in the operant devices, we measured the degree to which each algorithm produced the same triplet decision as did the birds. We focussed our evaluation on a held-out set of unambiguous sounds. Due to the inherent variability of behavioural decision data, the gold standard is not 100% but is an accuracy rate matching that of the birds. This rate cannot be measured directly since the ground truth is unknown for the test probes; we estimated it from their cycle accuracy as *estimated maximum attainable accuracy (cycle accuracy)*. We also calculated an upper bound based on the birds’ self- and inter-rater agreement on the test probes. This is an optimistic upper bound since it relies on a consistency assumption (namely, that each bird’s majority decision is the correct decision for a AXB triplet) as *estimated maximum attainable accuracy (consistency assumption).*

**Table 1:** Table of attributes for all learnt embedding space through our deep learning model.

|  | Emb-U | Emb-LU | Emb-LUA | Emb-Pre | Emb-PreUA | Emb-PreLUA |
| --- | --- | --- | --- | --- | --- | --- |
| pretrained (Pre) | - | - | - | X | X | X |
| Luscinia triplets (L) | - | X | X | - | - | X |
| unambiguous bird decisions (U) | X | X | X | - | X | X |
| ambiguous bird decisions (A) | - | - | X | - | X | X |

In the table, *pretrained (Pre)* refers to models that used Luscinia-U triplets to pre-train with and initialise the weights of the deep learning model; *Luscinia triplets (L)* refers to models that used Luscinia-U triplets for training; *unambiguous bird decisions (U)* refers to models that used the triplet decisions created by the bird judgements through our operant experiments; *ambiguous bird decisions (A)* refers to models that used the triplet decisions created by the bird judgements through our operant experiments. Note that some models used more than one source of triplets to train.

#### Luscinia parameter tuning

Separately, we used the bird decisions (from the unambiguous triplets) to tune the parameters of the Luscinia software (Lachlan, 2020). We refer to this tuned software as Luscinia-U.

Luscinia uses dynamic time warping (DTW) to align syllables, based on the trajectories of several acoustic features (Lachlan, 2020). In this case, these features were: fundamental frequency, peak frequency and mean frequency (log-transformed), fundamental and peak frequency change (the arcsin transform of the slope of these features on the spectrogram), normalized fundamental frequency (where the syllable-wide mean of fundamental frequency is subtracted from the fundamental frequency), Wiener entropy and harmonicity (measures of spectral complexity), and vibrato amplitude (a measure of sinusoidal signal in the fundamental frequency). Finally, time itself is included as a feature in order to penalize time warping. When two syllables are compared using this DTW process, a Euclidean distance is calculated over each of these features for each point in one syllable compared with each point in the other syllable. A dynamic algorithm then searches for an efficient alignment between the two syllables using these distances, and an overall dissimilarity is then calculated by averaging dissimilarities over the alignment.

A key challenge is to decide how to weight the sound features relative to each other. We developed a data-driven approach that used the same training data that we used to train neural networks. For each triplet in the dataset, we calculated the likelihood that the DTW algorithm would make the same choice as the birds. We then integrated log-likelihoods over the entire training dataset. We then used a Monte Carlo Markov Chain approach to find weightings that would maximise likelihood. The MCMC chain was initiated with equal weightings for each feature. We ran it for 10,000 generations, and treated the first 1,000 generations as burn-in. New parameter values were sampled from a log-Gaussian distribution with standard deviation 0.1. In each generation, parameters were normalized to sum to 1. We then estimated the mean parameter weightings from the last 9,000 generations of the MCMC and used those in a DTW analysis to measure the dissimilarity between each pair of syllables in the dataset. This dissimilarity matrix was then used for further analysis and comparison of different methods. The weightings of acoustic features before and after training are shown in Table S1.

We evaluated Luscinia-U directly, but we also experimented with using triplet decisions from that system as additional training data for deep learning, to supplement the bird decisions dataset. For this, we used the tuned Luscinia-U to generate additional AXB triplet decisions. These were then used as additional data, either to pretrain the dep learning network or simply pooled with the natural bird decisions during the main training phase. For this we filtered the Luscinia-U triplets down to only the coarse-level (‘easy’) distinctions using a minimum distance threshold, to minimise the risk of contradicting birds’ judgments.

### Other software

We compared the performance of our deep learning methods with current software used for similarity measures: Raven (Center for Conservation Bioacoustics, 2019), Sound Analysis Pro (Tchernichovski *et al*., 2000) and Luscinia (Lachlan, 2020) (with and without tuning). For each software package we extracted the (dis)similarity matrix of the evaluation set and used this to determine their AXB decisions. From this we calculated their accuracy at predicting bird AXB decisions.

### Song features

It is hypothetically possible that our trained deep learning algorithm would rely entirely on just one or a few basic underlying acoustic features, such as fundamental frequency. To test whether this happened, we first calculated, using Luscinia, for each syllable in the data-set, the syllable length, as well as the mean, maximum, minimum, start and end value of each of the following spectral acoustic features: fundamental frequency, peak frequency, median frequency, fundamental frequency change and harmonicity (Holveck *et al*., 2008). We then carried out an multiple regression on distance matrices (MRM) analysis using the r package *ecodist* (Goslee & Urban, 2007), a permutation based regression technique, predicting the dissimilarity matrix derived from the output of the machine learning algorithm Emb-LUA.

We included six predictor dissimilarity matrices based on the Euclidean distance for each of the five spectral acoustic features plus syllable length (10,000 permutations). The frequency parameters and length were log transformed, and all measures were normalised by their standard deviation.

## Results

### Training and Test outcomes

Using RFID technology we were able to continuously train and test each bird individually within its home cage and social group, enabling us to collect sound assessments over long time periods and on a much larger scale than conventional operant experiments. We trained 23 birds in 4 aviaries with in total 99 sets of stimuli. Over a period of 11 months we presented the birds with 1,116,214 trials, for which we received 927,158 responses. Of these trials, 25,999 trials were probe trials from which we collected 22,048 sound assessments from the birds. For each bird and each testing period we calculated a cycle accuracy, which is the overall accuracy in its responses to the training stimuli during a specific test period. Out of 99 stimulus sets, for 46 sets the birds reached a cycle accuracy of 65% or higher (*mean* ± *SE* = 72.07% ± 0.63%).

To assess whether birds made choices we expected we compared the birds’ assessments of similarity with assessments made by Luscinia. Birds with a cycle accuracy of at least 65% were more likely to choose the perch corresponding to the most-similar probe according to Luscinia, than in a random scenario (10,000 permutations) (Figure 2): random choice 95% confidence interval adjusted rank difference: Left choice = [0.005, 0.03]; median adjusted rank difference birds = −0.06; right choice =[0.07, 0.08]; median adjusted rank difference birds = 0.10; N = 15,141 responses to probe-stimulus triplets; 10,000 permutations.

**Figure 2:**
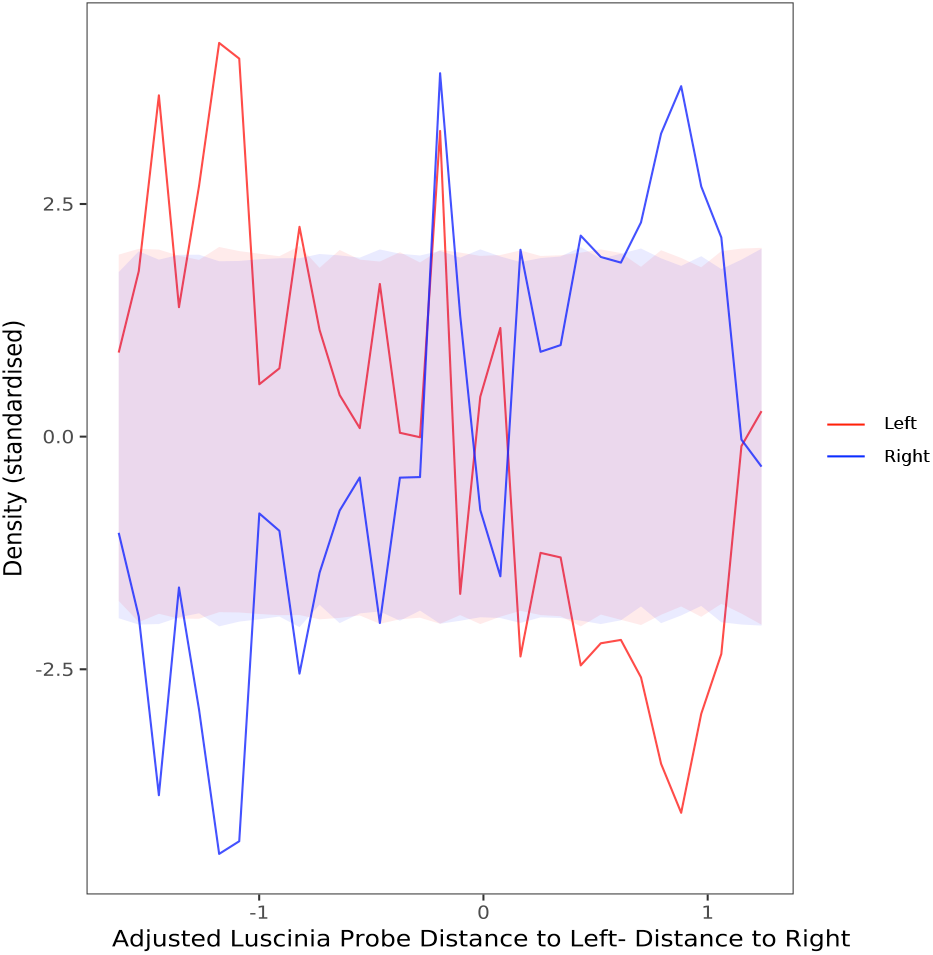
Birds’ choices in operant tests were more similar to computational methods than expected by chance. The figure shows the adjusted rank differences between probes and stimuli and the standardised density of choices for each of the sides of the apparatus. 95% confidence intervals of 1000 permutations of random responses are shown in light red and blue, and the birds’ actual assessments are shown in the dark red and dark blue lines. Random choice 95% confidence interval adjusted rank difference: Left choice = [0.005, 0.03]; median adjusted rank difference birds = −0.06; right choice =[0.07, 0.08]; median adjusted rank difference birds = 0.10

### Deep learning

To explore the sensitivity of our deep learning method to hyperparameter settings, we evaluated its performance at different dimensionalities of embedding space (Figure S3), and different ways of using bird decisions and Luscinia-U decisions in training (Table 1 and Figure 3). Good performance was obtained with embeddings of 16 dimensions or more, peaking at 64 dimensions (Figure S3).

**Figure 3:**
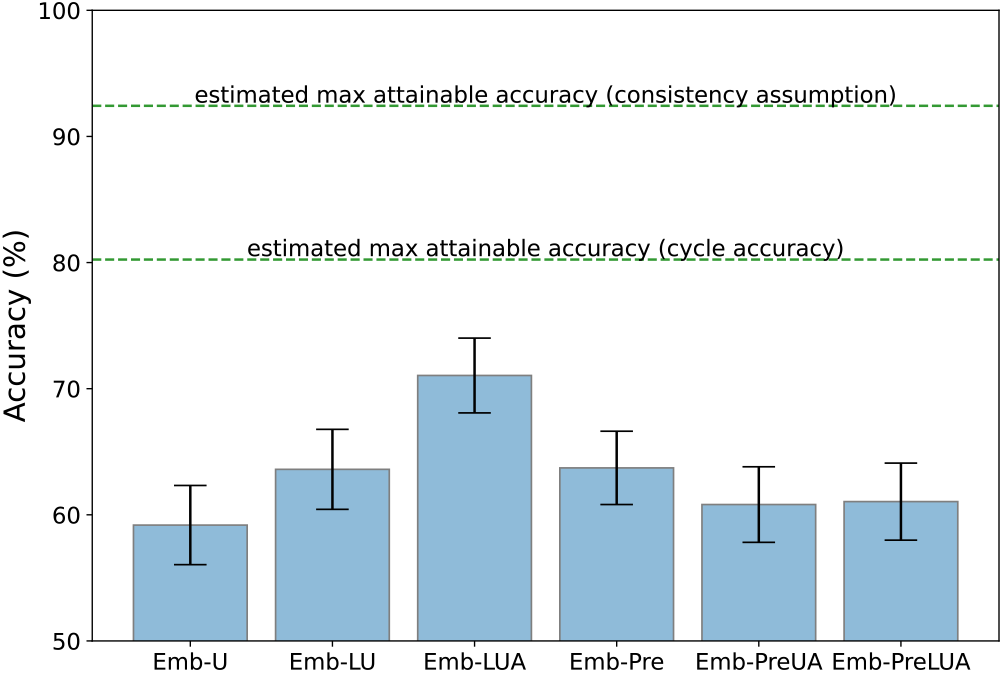
Deep learning performance is improved by our interventions, but not by pretraining. Accuracy performance of the different variants of our deep learning model, each model being described in Table 1. Error bars show 95% confidence intervals (bootstrap estimate). The upper lines indicate two different ways to estimate the maximum accuracy attainable in principle; 100% is not attainable since birds’ choices are not always consistent for the same stimulus set. An accuracy of 50% represents chance level.

We evaluated our algorithms by measuring the degree to which each algorithm produced the same triplet decisions as birds. We found that the deep learning performance is improved by our interventions, especially Emb-LUA—which used the unambiguous and ambiguous triplets as well as the additional Luscinia-U triplets—but not by pretraining (Figure 3). Using a mix of Luscinia-U decisions made on triplets of sounds along with perceptual decisions from the birds provided the best result.

We furthermore investigated the similarities between all models by projecting them in a 2-D space by using multi-dimensional scaling (MDS) (Borg *et al*., 2013) (Figure 5). In this visualisation, algorithms are closer together if they produce similar decisions, irrespective of whether those decisions are right or wrong. It illustrates that all three pretrained deep learning models remain very similar, and indeed very similar to Luscinia-U, the source of their pretraining, whereas our Emb-LUA model produces a different pattern of judgments.

**Figure 4:**
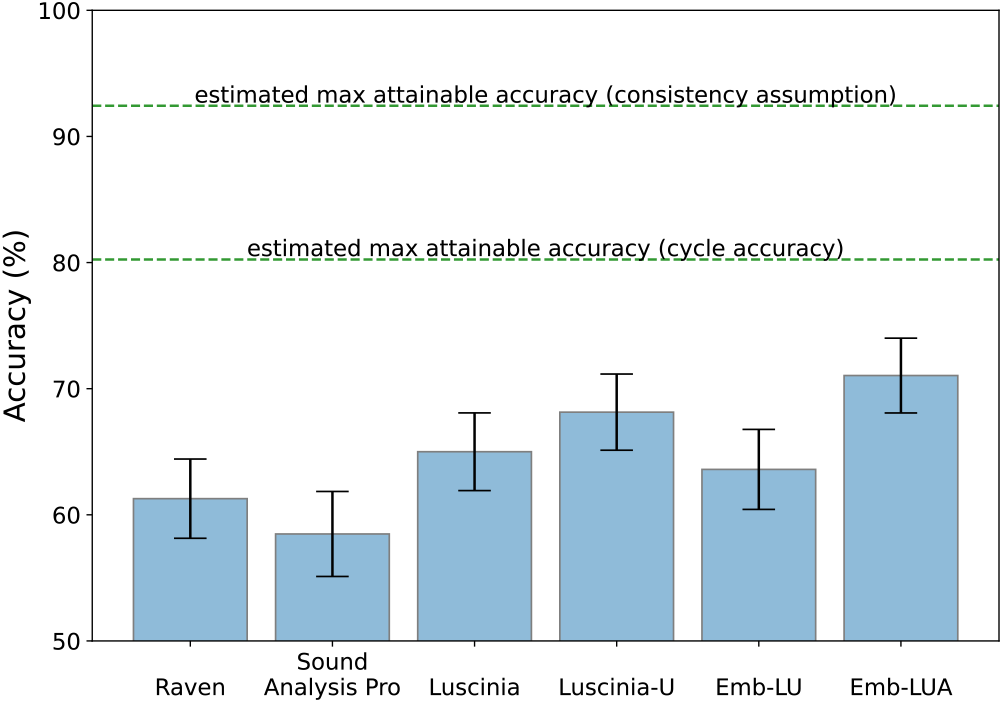
Accuracy of computational methods at predicting bird AXB decisions, evaluated on the evaluation set. Details are the same as in figure 4.

**Figure 5:**
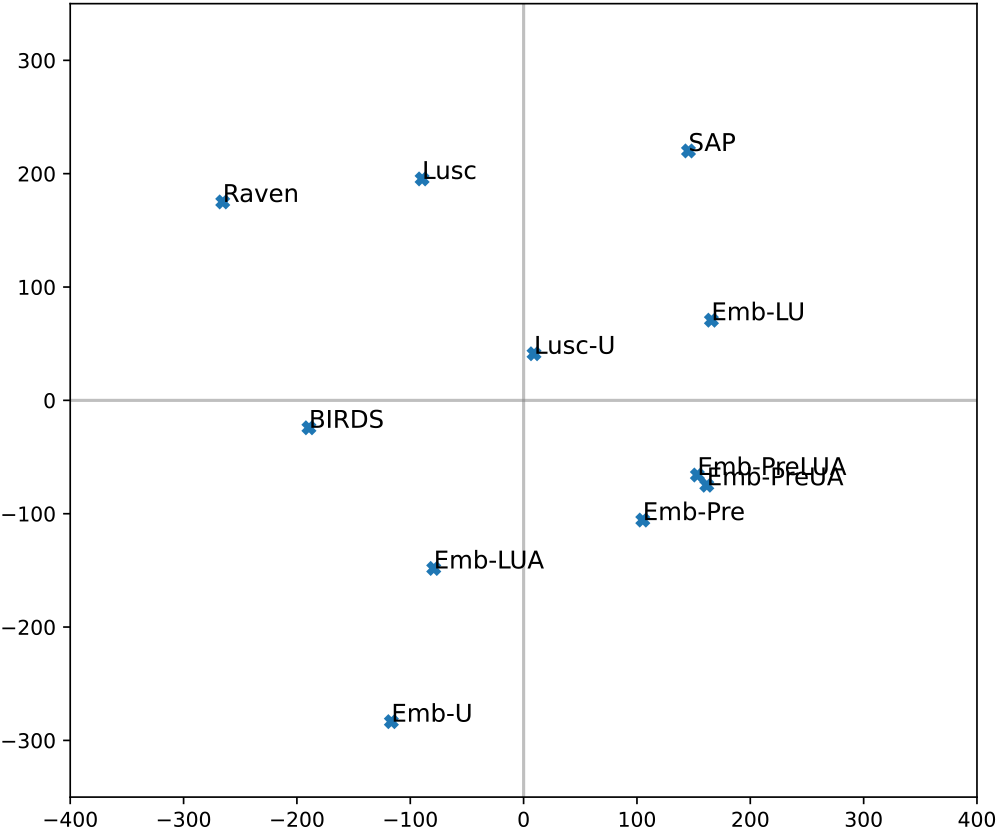
Different algorithms differ from birds’ judgements of similarity in different ways. 2-D multi-dimensional scaling (MDS) projection of the similarity between different algorithms based on their predictions. *BIRDS* is the ground truth provided by the bird decisions. Current software methods: *SAP* (Sound Analysis Pro). *Raven*, *Lusc* (Luscinia). *Lusc-U* represents the Luscinia software with tuned Luscinia parameters. Deep learning embedding models: *Emb-\** are as described in Table 1.

### Other software

When comparing the performance of our deep learning model with current software used for similarity measures—Raven (Center for Conservation Bioacoustics, 2019), Sound Analysis Pro (Tchernichovski *et al*., 2000) and Luscinia (Lachlan, 2020) (with and without tuning)—we found that deep learning Emb-LU trained using standard unambiguous triplets along with Luscinia-U triplets was unable to outperform these specialist tools (Figure 4). However, our deep learning model Emb-LUA trained on both unambiguous and ambiguous triplets along with Luscinia-U triplets can learn an embedding space that can reflect the birds’ perceptual judgements better than any current software publicly available. Comparison between Emb-LU and Emb-LUA leads us to conclude that even AXB triplets that yield inconsistent side decisions can provide useful information about perceived sound similarity, which helps to constrain the optimisation search space for deep learning. The difference in accuracy between the two best performing models, Emb-LUA and Luscinia-U, is 2.9 percentage points. We added all software methods to the 2-D space by using multi-dimensional scaling (MDS) (Figure 5).

When evaluating systems, we found that Sound Analysis Pro would yield similarities of precisely zero for some comparisons (in the case of high dissimilarity). When both pairs of similarities have zero values, we assume the three exemplars to be so dissimilar to each other that no similarity judgement can be computed from Sound Analysis Pro; this was the case for 103 triplets. To inspect whether these highly dissimilar triplets led to an unfair penalty for some systems, we repeated our evaluation with those 103 triplets excluded (Figure S4). This did lead to improved performance of Sound Analysis Pro, though the overall pattern of outcomes remained similar, with our Emb-LUA model maintaining the best performance over all other systems.

### Song features

The set of 26 acoustic features together explained 60.6% of variation in the machine learning dissimilarities (Figure S5). All features except fundamental frequency change were significant predictors of the machine learning dissimilarity matrix; fundamental frequency was the strongest predictor. A second analysis including just the single most informative acoustic feature, mean fundamental frequency, explained 56% of variation in the machine learning dissimilarities. Thus while a comprehensive set of acoustic features was, as expected, clearly related to the decisions that the machine learning algorithm made, much variation was left unexplained.

## Discussion

Making reliable comparisons of song recordings is essential for the field of bird song research. However, so far most comparison methods use HVA for validation, rather than the birds’ perception itself. We compiled a large corpus of zebra finches’ perceptual decisions of syllable similarities, which allowed us to validate existing methods, and also to train a deep neural network to assess song similarity.

Training and testing birds in a group setting allowed us to collect a dataset with over 22,000 probe assessments; to our knowledge no similar dataset has been generated and its size allows training of deep learning algorithms as we demonstrate. The social context of training, combined with relying on behavioural rather than neural responses potentially adds additional sources of noise compared with other approaches (e.g. Bell *et al*. (2015)). While in our experiments some birds reached accuracy levels of over 80%, some other birds’ performance remained consistently low in accuracy. One reason for these differences in accuracy may be that the group setting allowed some birds to scrounge others’ rewards (Giraldeau & Dubois, 2008), or to interfere with other individuals’ trials. In future experiments our method could be refined by adding an extra RFID antenna to the choice perches to limit choice registration to only the bird that started the trial. But even in social isolation, however, similar operant conditioning designs do not lead to birds approaching 100% accuracy: birds continue to frequently explore unrewarded options. These sources of error are substantially higher than when human observers are asked to generate perceptual training data-sets.

The birds’ choices in operant tests were, however, much more similar to computational methods than expected by chance. This suggests that overall the birds make decisions that correspond to acoustic characteristics. The differences between computational assessments and the birds’ decisions is difficult to interpret: we might expect that it can be explained partly by the noise in the data due to the birds’ behavioural variation in responses, and partly by differences between the algorithm and birds underlying perception. We have made the assumption that, taking into account our large sample size, our inclusion only of birds that performed to a reasonably high level on training choices, and evidence of agreement between trials when making the same choice on probes, that higher agreement between birds’ choices and computational judgement reflects a greater similarity between bird perception and the algorithm.

Using a mix of Luscinia-U decisions made on triplets of sounds along with perceptual decisions from the birds provided the best correspondence to bird decisions (Figure 3). We noticed that a pretraining phase using Luscinia-U triplet decisions led to worse performance. This finding is in contrast to much recent work in deep learning that makes heavy use of pretraining, especially when target datasets are small (Kong *et al*., 2020; Lasseck, 2018). It can be attributed to the fact that the pretrained space (driven by Luscinia’s similarity algorithm) is not appropriate, because it initialises the model such that its fine judgments follow Luscinia rather than bird judgments. Our alternative successful approach is to use both data sources simultaneously in the main training phase of deep learning.

The new algorithm cannot be explained well by standard acoustic features of the syllables. Several features together explained only around 60% of the variation in the machine learning dissimilarities. Moreover the relative contribution of different song features are similar to the parameter weight settings that resulted from tuning Luscinia with the birds’ decisions (Table S1). Although this wasn’t the primary goal of our study, these two sources of data also provide an indication of how birds integrate different perceptual features when assessing songs. It appears that mean fundamental frequency plays a particularly important role to zebra finches and that other temporal aspects of the pitch trajectory are, perhaps surprisingly, less critical. The birds (and deep learning algorithms) make use of spectro-temporal differences that are not reflected in standard acoustic features.

The two algorithms that had been trained with our training data set (Luscinia-U and our new machine learning algorithm) outperformed other methods of song comparison on our testing data-set. This supports the idea that data-informed methods of comparison may be more reliable than ones that rely solely on human intuition. Of the remaining algorithms that we tested, the untrained version of Luscinia’s dynamic time warping algorithm outperformed Sound Analysis Pro and the implementation of cross-correlation in Raven. Aspects of our study system and task may go some way to explaining these differences. Zebra finch song is highly spectrally complex compared to many birds’ song. Spectrographic cross correlation was originally employed on swamp sparrow song, where all spectral energy is concentrated at one frequency at any one time point, and it would seem logical that it might be harder to interpret spectrographic overlap in more complex scenarios. The algorithm used by Sound Analysis Pro was similarly designed for a particular task: to assess similarity between shared song syllables - i.e. pairs of syllables that share some degree of similarity. It was not designed to measure similarity between all pairs of notes, and beyond a certain level of dissimilarity, a floor of 0 is reached. This directly impacted its performance in our test: when removing affected triplets, its performance did not exceed that of Luscinia and our machine learning algorithm, but the three were relatively close in their performance. Most research on zebra finch song over the last two decades has used Sound Analysis Pro and our results would suggest that for tasks involving the assessment of the precision of song learning among relatively similar groups of syllables, these analyses have a reasonable degree of validity.

Although the algorithm we introduced here outperforms other commonly used methods and presents a major step forward in the measurement of song similarity, some care has to be taken. In this experiment we have presented birds with single song syllables. Although these isolated syllables, rather than full song, may have sounded less natural to the birds, and also limited the possibilities of measuring higher levels of song perceptions, they made it possible to study sound perception using sounds with features they are naturally attuned to. Furthermore, although all syllables used originated from a single population (different from the experimental population), since the variation in syllables within a zebra finch colony is large compared to that between colonies (Lachlan *et al*., 2016) (but see Wang *et al*. (2022)), we do not expect that this limits the generalisability of the algorithm for different zebra finch colonies. Finally, the current algorithm is only based on zebra finch perceptual judgements which may not be the same for all bird species. Different species have different hearing sensitivities, and may differ in how they weight different sound features (Dooling & Prior, 2017b; Dooling *et al*., 1992; Okanoya & Dooling, 1988). Collecting similar perceptual judgements from a range of other species will enable us to compare differences in perception and will improve our overall ability to accurately measure song similarity.

## Author contributions

LZ, VM, DFC, DS, RFL designed the research; LZ, VM, JG, DS, RFL performed the research; LZ, VM, DS, RFL analysed data; LZ, VM, DS, RFL wrote the paper. All authors contributed critically to the drafts and gave final approval for publication.

## Acknowledgements

This research was supported by BBSRC research Grant. No. BB/R008736/1 “Machine Learning for Bird Song Learning.”

## Data accessibility

Both the code and the data will be stored on Zenodo

## Notes

https://doi.org/10.5281/zenodo.5545872

### Competing Interest Statement

The authors have declared no competing interest.

