## Supplementary Information for "Bird song comparison using deep learning trained from avian perceptual judgments"

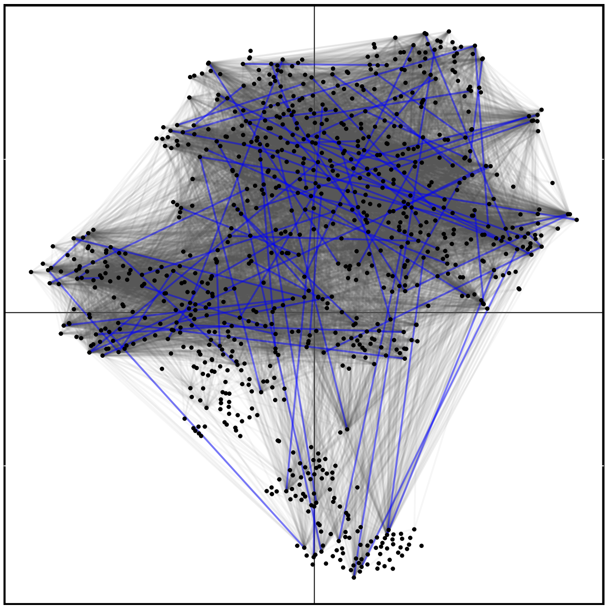

Figure S1: Training stimuli and probes projected in 2 dimensional space (extracted from 64 dimensional embedding using T-SNE for the projection with euclidean distances). Sets of training stimuli are connected with blue lines, connections between probes and training stimuli with grey lines.

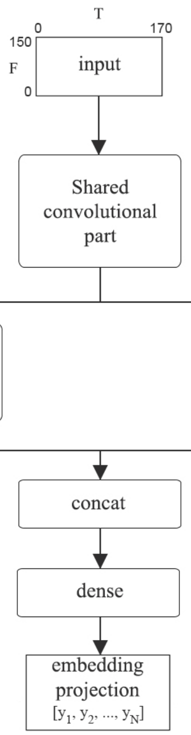


Figure S2: Overview of deep learning model architecture.

Table S1: Table of parameter settings for Luscinia.

| Parameter | Weighting prior to training | Weighting post-training |
|---|---|---|
| Mean frequency | 0.0925 | 0.0568 |
| Peak frequency | 0.0925 | 0.0273 |
| Fundamental frequency | 0.0925 | 0.3788 |
| Peak frequency change | 0.0925 | 0.0122 |
| Fundamental frequency change | 0.0925 | 0.0597 |
| Normalized fundamental frequency | 0.0925 | 0.2369 |
| Wiener entropy | 0.0925 | 0.00021 |
| Harmonicity | 0.0925 | 0.0832 |
| Time | 0.25 | 0.0443 |

The table shows the parameter weightings in Luscinia set before training, and achieved after training. Luscinia's DTW algorithm aligns acoustic features as they vary over the length of the syllable. Different features can be weighted more or less heavily.

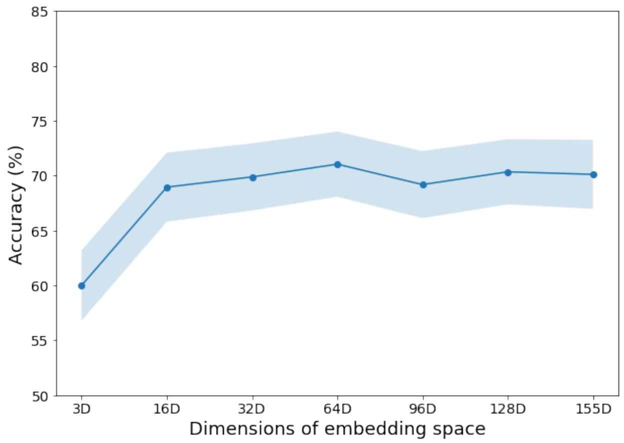


Figure S3: Accuracy (%) of deep learning models based on the learnt embedding space dimensions.

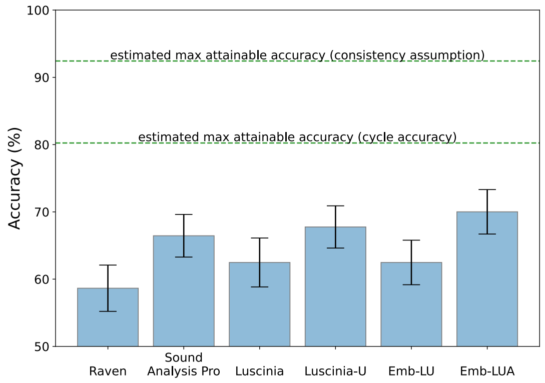


Figure S4: Accuracy (%) results on evaluation set after removing triplets that produce zero '%similarity' values in Sound Analysis Pro for both pairs in a triplet. Details are the same as in figure 4.

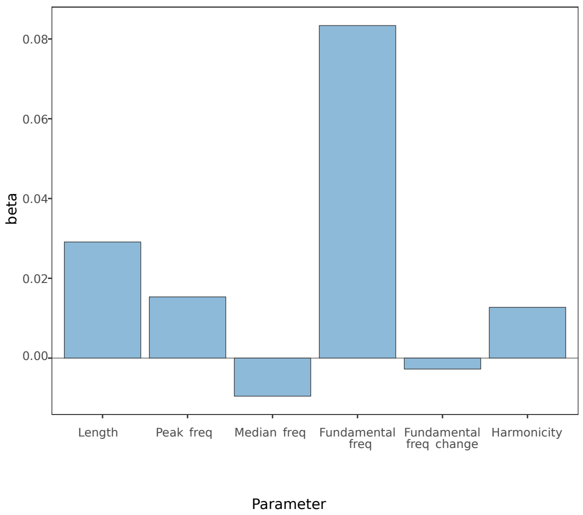


Figure S5: Acoustic predictors of machine learning. Each bar represents dissimilarity for the measures taken from fundamental frequency.
